# End-to-end deep learning approach to mouse behavior classification from cortex-wide calcium imaging

**DOI:** 10.1101/2023.04.05.535664

**Authors:** Takehiro Ajioka, Nobuhiro Nakai, Okito Yamashita, Toru Takumi

**Affiliations:** Department of Physiology and Cell Biology, Kobe University School of Medicine, Chuo, Kobe 650-0017, Japan; Department of Computational Brain Imaging, ATR Neural Information Analysis Laboratories, Seika, Kyoto 619-0288, Japan; RIKEN Center for Biosystems Dynamics Research, Chuo, Kobe 650-0047, Japan

## Abstract

Deep learning is a powerful tool for neural decoding, broadly applied to systems neuroscience and clinical studies. Interpretable and transparent models which can explain neural decoding for intended behaviors are crucial to identify essential features of deep learning decoders in brain activity. In this study, we examine the performance of deep learning to classify mouse behavioral states from mesoscopic cortex-wide calcium imaging data. Our convolutional neural network (CNN)-based end-to-end decoder combined with recurrent neural network (RNN) classifies the behavioral states with high accuracy and robustness to individual differences on temporal scales of sub-seconds. Using the CNN-RNN decoder, we identify that the forelimb and hindlimb areas in the somatosensory cortex significantly contribute to behavioral classification. Our findings imply that the end-to-end approach has the potential to be an interpretable deep learning method with unbiased visualization of critical brain regions.

**Author Summary:** Deep learning is used in neuroscience, and it has become possible to classify and predict behavior from massive data of neural signals from animals, including humans. However, little is known about how deep learning discriminates the features of neural signals. In this study, we perform behavioral classification from calcium imaging data of the mouse cortex and investigate brain regions important for the classification. By the end-to-end approach, an unbiased method without data pre-processing, we clarify that information on the somatosensory areas in the cortex is important for distinguishing between resting and moving states in mice. This study will contribute to the development of interpretable deep-learning technology.

## Introduction

Neural decoding is a method to understand how neural activity relates to perception systems and the intended behaviors of animals. Deep learning is a powerful tool for accurately decoding movement, speech, and vision from neural signals from the brain and for neuroengineering such as brain-computer interface (BCI) technology that utilizes the correspondence relationship between neural signals and their intentional behavioral expressions (Craik et al., 2019; LeCun et al., 2015; Livezey and Glaser, 2021). In clinical studies, electrical potentials measured by implanted electrodes in a specific brain area, such as the motor cortex, were often used to decode the intended movements such as finger motion, hand gesture, and limb-reaching behavior (Hochberg et al., 2012; Pan et al., 2018; Schwemmer et al., 2018; Skomrock et al., 2018). In contrast, neural decoding of the movements with whole-body motion, such as running and walking, remains uncertain due to measurements of neural activity from the entire brain under immobilized conditions in functional magnetic resonance imaging (fMRI) and magnetoencephalography (MEG) scanners and contamination of noise signals (e.g., non-neuronal electrical signals during muscular contraction) in electroencephalography (EEG) recording. It is challenging to decode voluntary behaviors from brain dynamics that contain complex information processing from motor planning to sensory feedback during the execution of a movement.

The calcium imaging technique allows us to measure *in vivo* neural activity during behavioral conditions from microscopic cellular to mesoscopic cortex-wide scales (Ren and Komiyama, 2021). Recent studies suggest that cellular activities have enough resolution for decoding behaviors. The cellular imaging data using microendoscopy in the hippocampal formation was used to decode free-moving mouse behaviors (Chang et al., 2021; Etter et al., 2020; Murano et al., 2022) by a Baysian- and a recurrent neural network (RNN)-based decoders. In addition, a convolutional neural network (CNN) is also used to predict the outcome of lever movements from microscopic images of the motor cortex in mice (Li et al., 2019). On the other hand, it is little known whether mesoscopic cortex-wide calcium imaging that contains neural activity at the regional population-but not the cellular resolution is applicable for neural decoding of animal behaviors. This mesoscopic strategy may be appropriate for end-to-end analyses since it deals with substantial spatiotemporal information of neural activity over the cortex.

Minimal preprocessing of input data can attenuate arbitrary interference for neural decoding. CNN is most applicable to image data, while RNN is often used for sequential inputs, including time-variable data (LeCun et al., 2015). By taking advantage of these architectures, we developed a two-step CNN-RNN model for decoding behavioral states from the mesoscopic cortical fluorescent images without intermediate processing. Moreover, it is desired to identify biologically essential features for deep learning classification to make the models interpretable and transparent for explanations of neural decoding as suggested by XAI-Explainable Artificial Intelligence (Gunning et al., 2019). To this end, we developed a visualization method of the features that contributed to the performance of the CNN-RNN-based classifications and identified the somatosensory areas are the most significant features for the type of behavioral states during voluntary locomotion behavior. This unbiased identification was supported by separate analyses of regional cortical activity using deep learning with RNN and the assessment by Deep SHAP, a developed Shapley additive explanations (SHAP) for deep learning (Lundberg and Lee, 2017; Vega García and Aznarte, 2020). Our findings demonstrate possibilities for neural decoding of voluntary behaviors with the whole-body motion from the cortex-wide images and advantages for identifying essential features of the decoders.

## Results

To perform behavior classification from the cortical activity with deep learning, we used the previously reported data composed of mesoscopic cortex-wide calcium imaging in the mouse, which exhibits voluntary locomotion behavior in a virtual environment under head-fixed conditions (Nakai et al., 2023). The fluorescent calcium signals of the cortex were imaged at a frame rate of 30 frames/s during a 10-min session (18,000 frames/session) from behaving mice (**Figs 1A–1B**). Two behavioral states (run or rest) were defined by a threshold of the speed of locomotion (>0.5 cm/s) and binarized as 1 for a run and 0 for rest in each frame. The proportion of run state differed according to individual mice (mean ± SD; mouse ID1, 36 ± 8 % (n = 11 sessions); ID2, 66 ± 22 % (n = 12 sessions); ID3, 65 ± 16 % (n = 14 sessions); ID4, 58 ± 11 % (n = 15 sessions); ID5, 80 ± 8 % (n = 12 sessions); **Fig 1C**). To generalize decoding across individuals, we assigned the data to training, validation, and testing at the ratio of 3:1:1 on a per-mouse basis (**Fig 1D**). Thus, we generated 20 models for all combinations and classified the test data with each.

**Fig 1.**
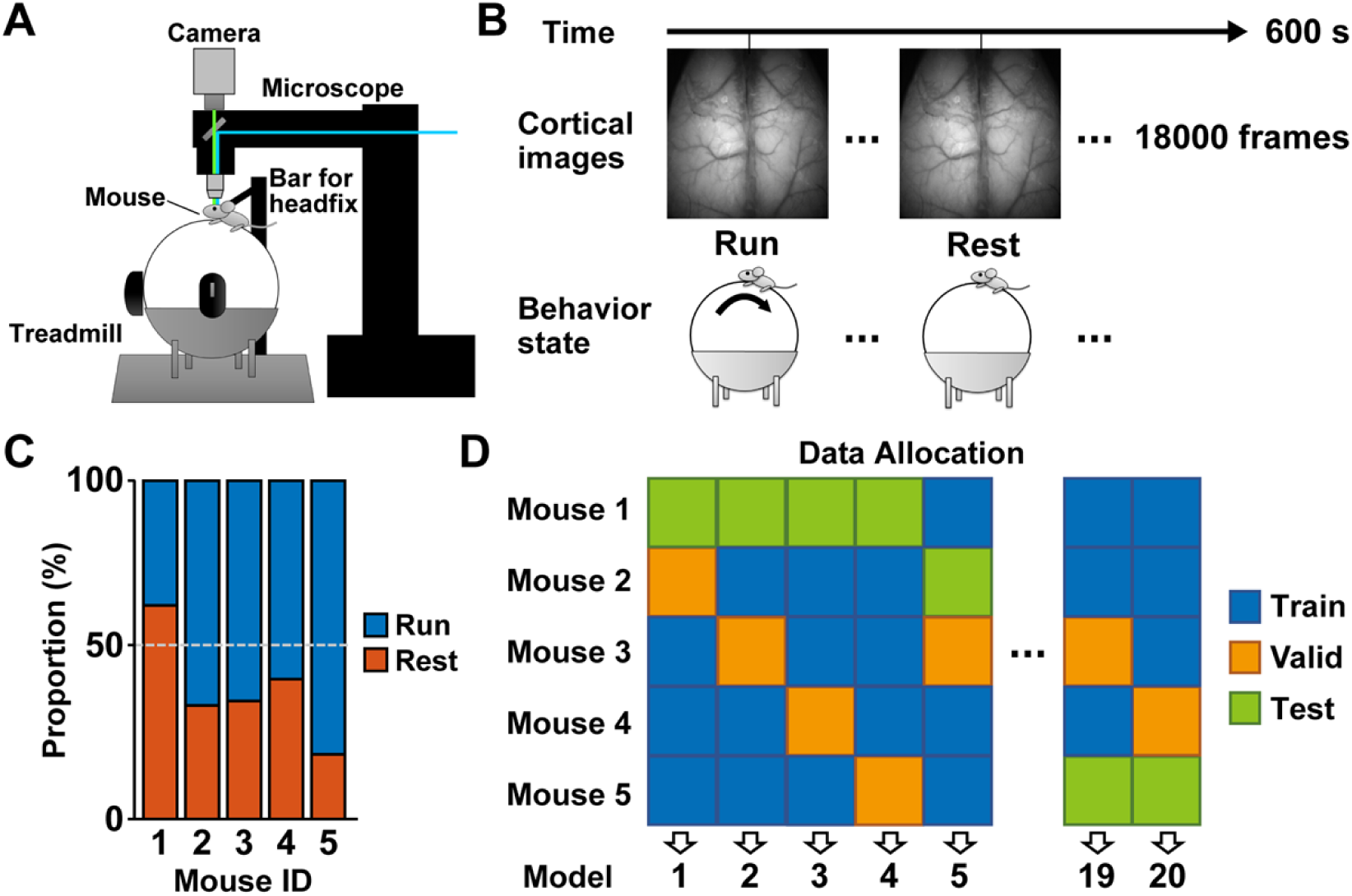
Cortical activity and behavioral states in behaving mice. (A) A schematic illustration of the experimental setup for measuring mesoscopic cortical calcium imaging and locomotor activity. (B) Images were obtained at 30 frames per second during a 600 s session. The label of behavioral state was based on locomotion speed (>0.5 cm/s) at the corresponding frame. (C) Proportions of the behavioral states in each mouse (n = 11–14 sessions from 5 mice). (D) The data allocation on a per-mouse basis. The data of each mouse was assigned at the ratio of 3:1:1 for training (Train), validation (Valid), and testing (Test).

### CNN-based end-to-end deep learning accurately classified behavioral states from functional cortical imaging signals

We tried to classify the behavioral states from images of cortical fluorescent signals using deep learning with CNN. To handle the single-channel images obtained from calcium imaging, we converted a consequence three images into a pseudo-3-channel RGB image by combining the previous and next images with the target image (**Fig 2A**). First, we trained CNN with EfficientNet B0 (Tan and Le, 2020), where the individual RGB images were used for input data. The binary behavior labels were used for output (**Fig 2B**). We used the pre-trained model on ImageNet for the initial weight values in training. In training, the loss was reduced by increasing epochs in CNN decoders (**Fig 2D, left**). However, in validation, the loss was increased every epoch (**Fig 2D, left**), suggesting that models fell into overlearning during CNN training. We chose a model with the lowest loss in the validation as a decoder at each data allocation. The decoder’s performance was evaluated by the area under the receiver operating characteristic curve (AUC) for all test data frames. The decoder using CNN alone classified the behavioral states with about 90% accuracy (0.896 ± 0.071, mean ± SD, n = 20 models; **Fig 2E**).

**Fig 2.**
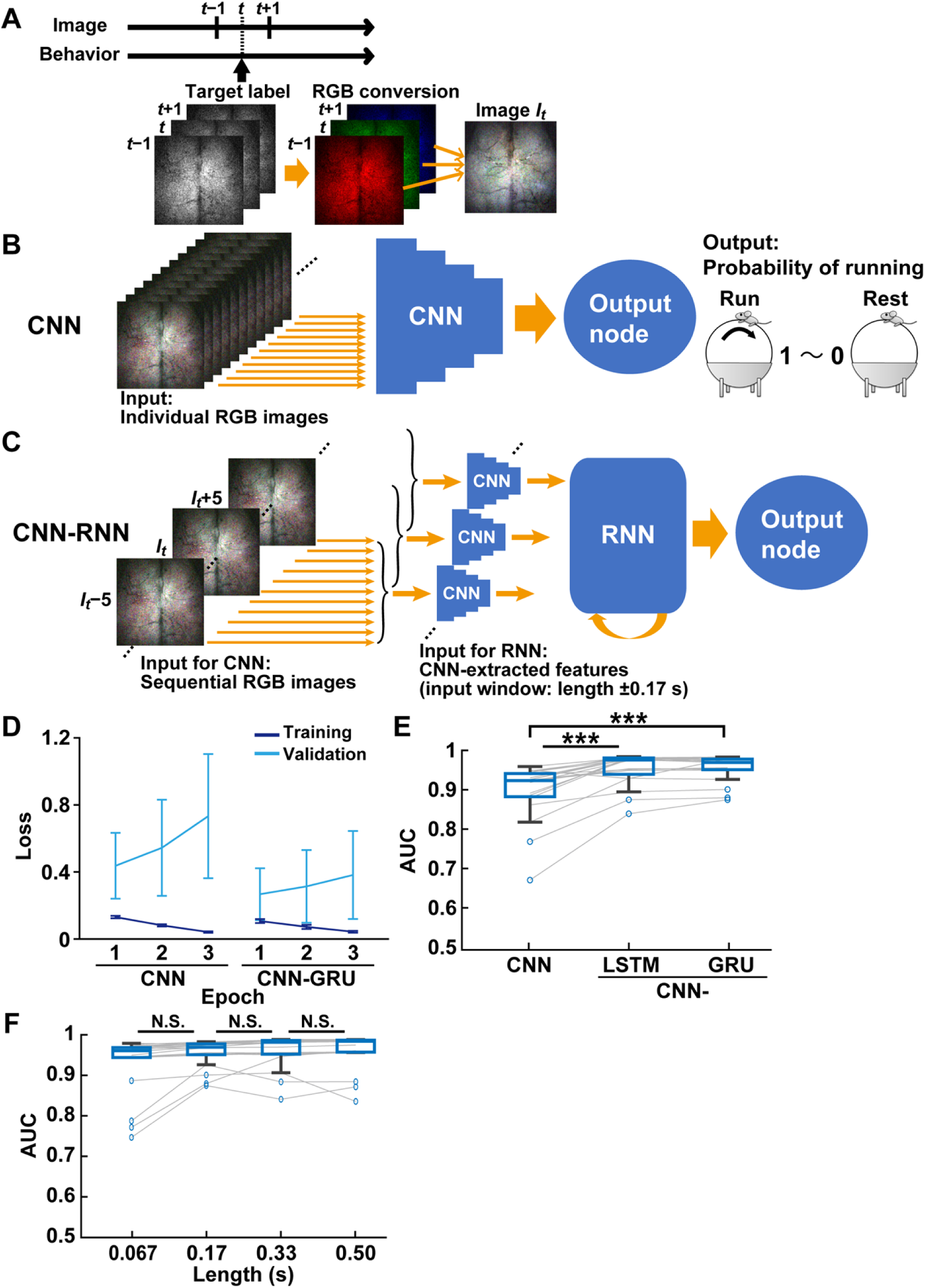
Behavioral state classification using deep learning with CNN. (A) Image preprocessing for deep learning with CNN. An image at frame *t* with images at neighboring frames (frame *t* −1 and *t* +1) was converted to an RGB image (image *I_t_*) labeled with the behavioral state. (B) Schematic diagram of the CNN decoder. CNN was trained with individual RGB images. Then, CNN outputs the probability of running computing from the 1,820 extracted features for each image. (C) Schematic diagram of the CNN-RNN decoder. The pre-trained CNN extracted 1,820 features from individual RGB images in the first step. In the second step, a series of 1,820 extracted features obtained from consecutive images (e.g., eleven images from *I_t_* −5 to *I_t_*+5 (= input window, length ±0.17 s)) were input to GRU-based RNN. Then, the RNN output probability of running. (D) Loss of CNN and CNN-GRU during training and validation across three epochs. (E) The area under the receiver operating characteristic curves (AUC) was used to indicate the accuracy of decoders. The performance of decoders with CNN, CNN-LSTM, and CNN-GRU. \*\*\**P* < 0.001, Wilcoxon rank-sum test with Holm correction, n = 20 models. (F) The performance of CNN-GRU decoders was not significantly different between different lengths of the input window. N.S., not significant, Wilcoxon rank-sum test with Holm correction, n = 20 models.

To improve the performance of decoding, we then created a two-step deep learning architecture that combines CNN with long short-term memory-(LSTM) (Hochreiter and Schmidhuber, 1997) or gated recurrent unit-(GRU) (Cho et al., 2014) based RNN, in which the output at the final layer of the CNN was compressed by average pooling and connected to the RNN (**Fig 2C**). In this stage, input data was the sequential RGB images from −0.17 s to 0.17 s from the image *t,* located at the center of the input time window. We used weights of the former CNN decoders for setting the initial values in two-step CNN-RNN. As with CNN decoders, the loss of two-step CNN-RNNs was reduced by the increment of epochs in training, whereas it was increased in validation (**Fig 2D, right**). The performance of behavior state classification was upgraded using two-step CNN-RNNs regardless of individual cortical images and behavioral activities (GRU, 0.955 ± 0.034; LSTM, 0.952 ± 0.041; mean ± SD, n = 20 models; **Fig 2E**). In addition, we confirmed that the classification accuracy was not significantly affected by the length of the input window ranged from 0.067 s to 0.50 s in the two-step deep learning (**Fig 2F**). These results demonstrate that deep learning decoding with CNN classifies locomotion and rest states accurately from functional cortical imaging consistently across individual mice, and the performance of the decoding can be improved by combining it with RNN.

### The somatosensory area contains valuable information on the behavioral classification

To make deep learning decoding interpretable, we tried to quantify the critical areas of images which contributed to the behavioral classification in the CNN-RNN decoder. We calculated and visualized the importance score in subdivisions of images in each decoder using a newly developed method named cut-out importance (see Methods for details). Briefly, a subdivision of the image was covered with a mask filled with 0 before training. The decoder trained with the masked images was compared with the decoder with original unmasked images (**Fig 3A**). The importance score indicates how much the decoder’s performance was affected by the masked area. As a result, the highest importance score was detected slightly above the middle of the left hemisphere (0.054 ± 0.045; mean ± SD, n = 20 models; **Fig 3B**). The symmetrical opposite area is also higher than other subdivisions within the right hemisphere (0.024 ± 0.014). This laterality seemed to be derived from individual differences (**S1 Fig**). These subdivisions corresponded to the anterior forelimb and hindlimb areas of the somatosensory cortex (**Fig 3C**; **S2 Fig**).

**Fig 3.**
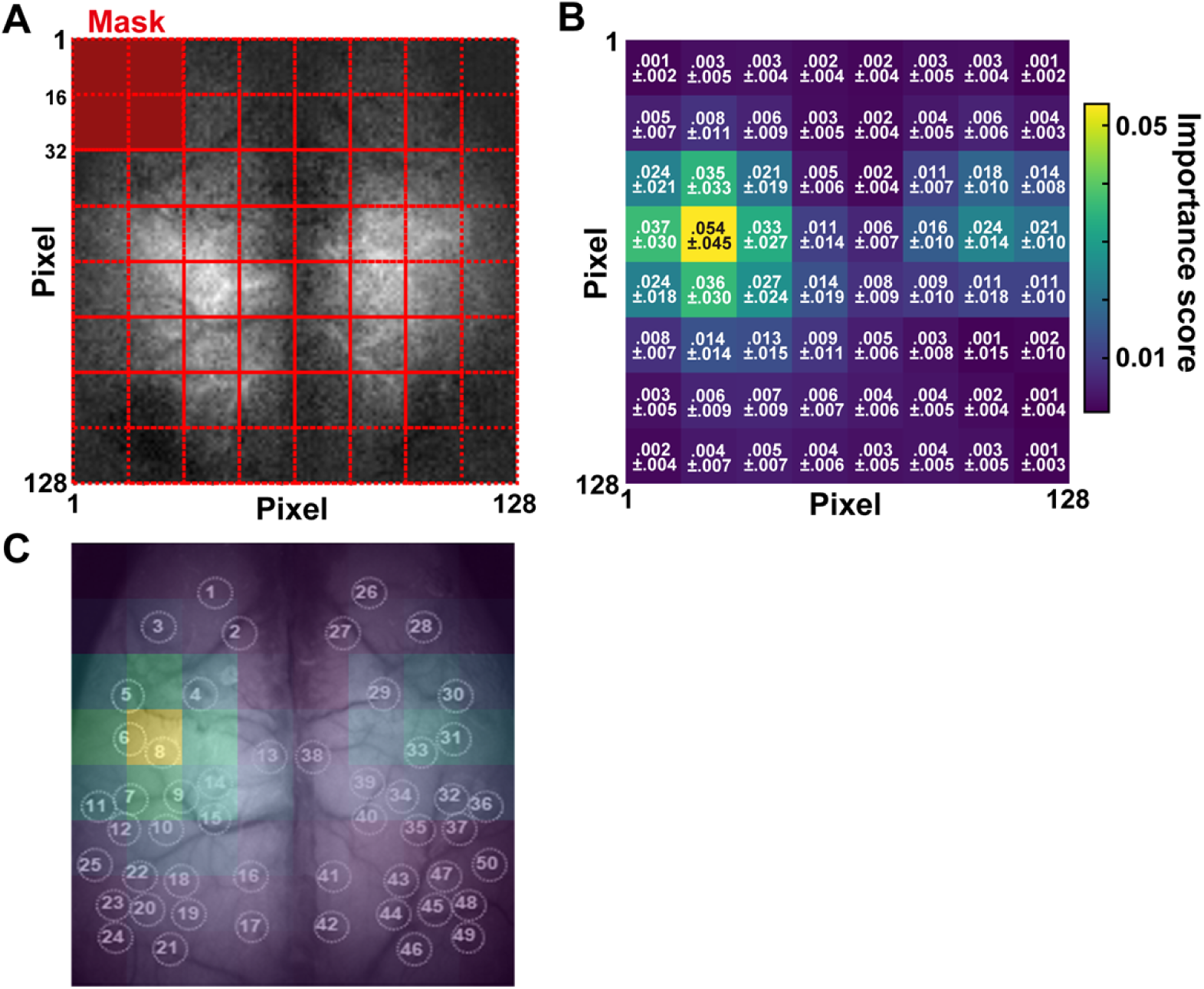
Visualization of essential features in CNN-RNN decoder. (A) An importance score was calculated by averaging differences from classification accuracy using a 1/16 masking area in each image (see Methods for details). (B) Importance scores in each subdivision (mean ± SD, n = 20 models). (C) Overlay of importance scores on the cortical image with ROI positions. See S2 Fig for ROIs 1–50.

### Regional cortical activity is applicable for the behavioral classification using RNN **decoders**

To confirm the contribution of the somatosensory cortex in the decoding performance, we designed RNN decoders to classify the behavioral states from activities of the specific cortical areas. For this purpose, the fluorescent signals at 50 regions of interest (ROIs) in the cortex were analyzed as regional cortical activities that accord with known cortical parcellations of the mouse brain (**S2 Fig**; (Nakai et al., 2023)). To reduce baseline fluctuation of cortical activity, we performed data preprocessing by subtracting a 1,000-frame moving average from the normalized fluorescent signals at each ROI (**S3 Fig**).

At the beginning of the deep learning decoding with RNN, we used a GRU architecture and set an input window of size 31, including a one-second duration of cortical activity that ranged from −0.5 s (−15 frames) to 0.5 s (+15 frames) from the behavioral state-target label (frame *t*) (**Fig 4A**). To train the deep learning models, we used the ±0.5 s input window with a one-frame sliding window for a total of 1,152,000 frames data (n = 64 sessions). The random batches of size 256 with Adam optimizer (https://keras.io/api/optimizers/adam/ (Kingma and Ba, 2017)) and binary cross-entropy loss function were used as model parameters. The models were trained across 30 epochs to converge the loss substantially. In the training data, the loss was reduced in the first 10 epochs, with a slight improvement in the following epochs, and the accuracy was dramatically improved and almost saturated within the first 10 epochs (**Fig 4B**). In the validation, although changes of loss and accuracy behaved similarly, the loss was about twice, and the accuracy was slightly decreased compared to the training (**Fig 4B**). We chose a model with the lowest loss in the validation as a decoder at each data allocation. Then, the decoders classified all frames of the test data into the two behavioral states in good agreement with the behavioral labels (**Fig 4C**), supported by the AUC (**Fig 4D**). The GRU decoder trained with preprocessing data (mean ± SD; GRU, 0.974 ± 0.014; n = 20 each; **Fig 4E**) showed significantly higher performance of behavioral classification with high accuracy than the GRU decoder trained with un-preprocessing data (Raw, 0.911 ± 0.057). Both performances were considerably higher than the control decoder, a null model trained with randomly assigned behavioral labels (Random, 0.492 ± 0.031).

**Fig 4.**
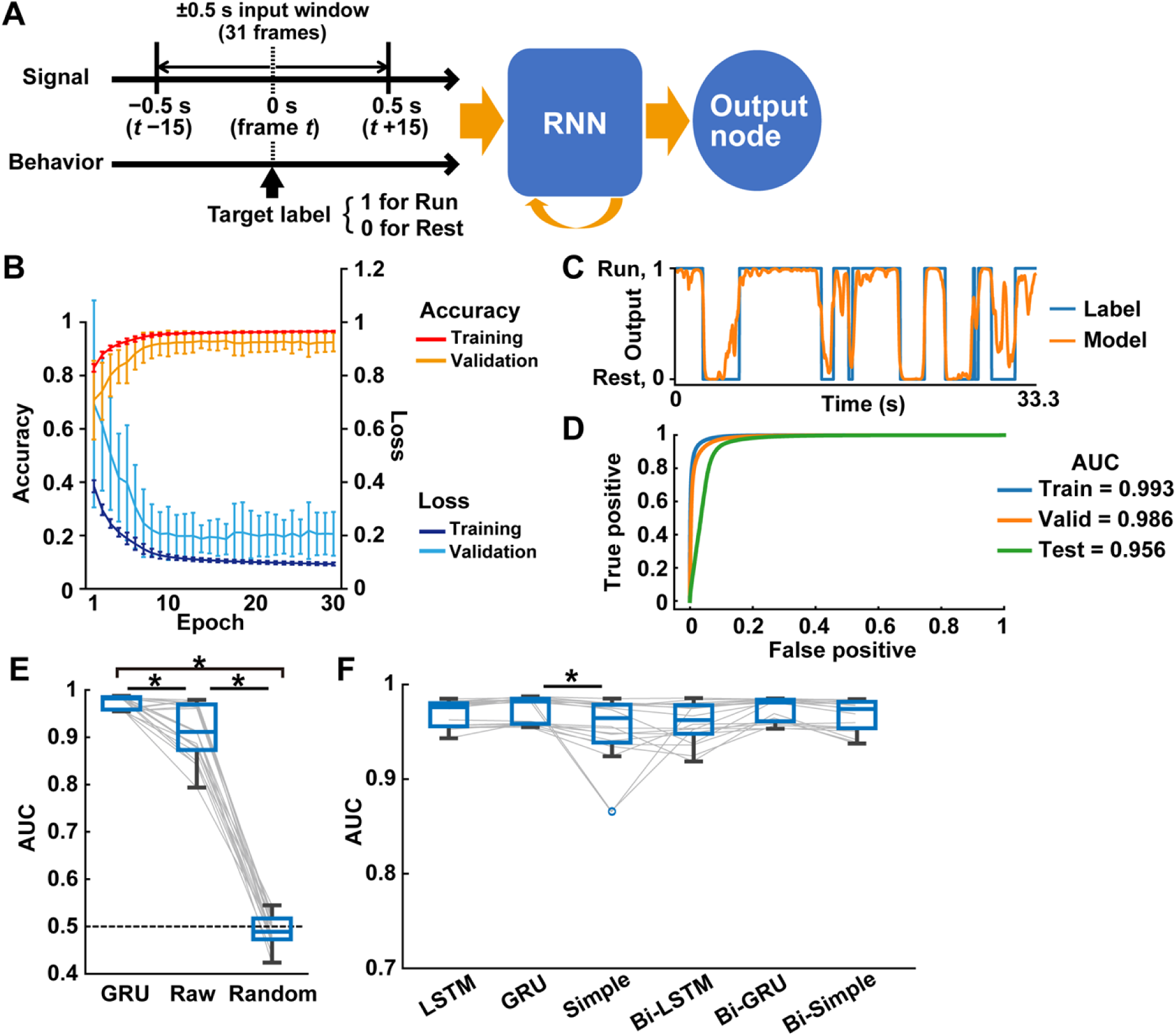
Behavioral state classification from cortical activity using deep learning with RNN. (A) Schematic overview of the RNN decoder for the behavioral state classification. Input is the cortical activities ranging from 0.5 s before (*t*−15 frames) to 0.5 s after (*t*+15 frames) the target frame *t,* which is labeled with a behavior state (1: run, 0: rest). The RNN decoder outputs the probability of behavioral states for all frames of testing data. (B–D) Example of the GRU decoder performance. (B) Learning curve during training and validation across 30 epochs. Loss indicates the cross entropy loss between the outputs and behavioral labels. Accuracy was the percentage of agreement with the label when the output was binarized at a 0.5 threshold. Mean ± SD, n = 20 models. (C) A trace of the output values of a representative decoder and actual behavioral labels in the first 33.3 s of testing data. (D) The receiver operating characteristic curves in the training, validation, and testing data. (E) The performance of GRU decoders trained with preprocessed data (GRU), non-preprocessed data (Raw), and randomly shuffled data (Random). \**P* < 0.05, Wilcoxon rank-sum test with Holm correction, n = 20 models. (F) The decoder performance using six types of RNN architectures. LSTM, GRU, simple RNN (Simple), and their bidirectional ones (Bi-). \**P* < 0.05, Wilcoxon rank-sum test with Holm correction, n = 20 models.

We next examined how much the architectures of RNN affect the decoder performance. All decoders classified behavioral states with high accuracy over 0.95 on average (mean ± SD; LSTM, 0.970 ± 0.013; Simple, 0.953 ± 0.035; Bi-LSTM, 0.960 ± 0.020; Bi-GRU, 0.974 ± 0.012; Bi-Simple, 0.967 ± 0.016; **Fig 4F**), while the simple RNN decoder only underperformed compared with the GRU decoder (*P*<0.05, Wilcoxon rank sum test with Holm correction). Given the accuracy and variance in these decoder performances, GRU and bidirectional GRU architectures are most suitable for the behavioral classification from cortical activity. We used, hereinafter, GRU but not bidirectional GRU as an RNN architecture to simplify the process and time of computing. We investigated whether the temporal specificity of the input data affects the performance of GRU decoders. The initial setting of the length of the input window was 0.5 s when the length contains information on cortical fluorescent signals ranging between 0.5 s before and after the center of the input window (i.e., 0 s). The shift 0 s was initially chosen, which means the position of the behavioral label at 0 s (**Fig 5A**). Regarding the analysis of length, the accuracy of the decoder performance from length 0.33 s to 1.0 s did not differ (**Fig 5B**). Only the accuracy was significantly decreased at length 0.17 s, suggesting that a temporally enough length (≥0.33 s) of input window is needed to obtain information of behavioral states from cortical activity. We then examined the temporal distance of the decoding target from the center of the input window by shifting the position of the target labels in a time range from −2 s (backward in time) to 2 s (forward in time) (**Fig 5C**). The accuracy of back-shifted target labels gradually but significantly decreased with distance from the center of the input window. Similarly, in the forward shift of target labels, the performance was significantly degraded when the target labels were set to more than 0.33 s distant from the center of the input window. These results suggest that our decoders are more fitting for predicting current states than future and past states of behaviors.

**Fig 5.**
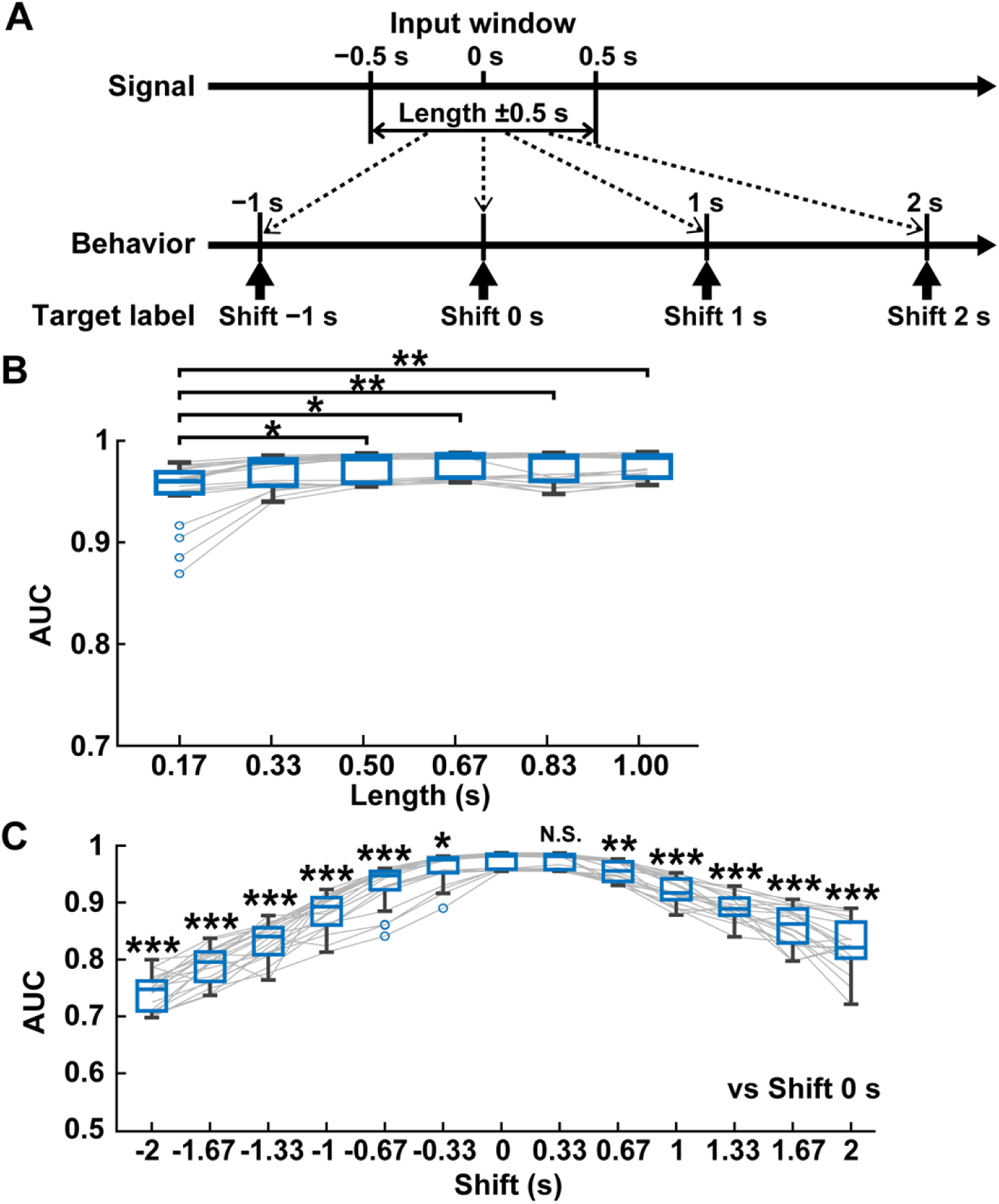
Comparison of input window length and target label’s temporal position. (A) Examples of input window and position of the target labels for behavior classification were shown. “Length” defines the duration of the input window, which ranges arbitral time (e.g., 0.5 s) before and after the center of the input window (0 s). “Shift” defines the temporal location of the target label of behavior classification from the center of the input window. The length 0.5 s and the shift 0 s were used for the criteria for evaluation. (B) The decoder performance of different lengths using a fixed shift 0 s. \**P* < 0.05, \*\**P* < 0.01, Wilcoxon rank-sum test with Holm correction, n = 20 models. (C) The decoder performance of different shifts using a fixed length of 0.5 s. N.S., not significant, \**P* < 0.05, \*\**P* < 0.01, \*\*\**P* < 0.001, Wilcoxon rank-sum test with Holm correction compared with shift 0 s, n = 20 models.

### Cortical activity in the somatosensory limb areas contributes to the behavioral classification

Finally, we assessed how much cortical areas significantly impact the GRU decoder using Deep SHAP (see Methods for details). We visualized a SHAP value which is the index to what extent each feature contributes to the behavioral classification in the trained models. The SHAP values in a model were calculated against each input window from ∼5% of randomly selected test data. The absolute SHAP values were averaged across all models to quantify the degree of importance in cortical areas (**Fig 6A**). The remarkably high SHAP values were detected in the anterior regions of the somatosensory forelimb (FLa, ROIs 6 and 31) and hindlimb (HLa, ROIs 8 and 33) areas. The peaks of SHAP values were observed around +0.1 s after the center of the input window. Although SHAP values of many cortical areas surpassed those in null models, overall, the magnitudes were smaller than the somatosensory areas (**Fig 6B**).

**Fig 6.**
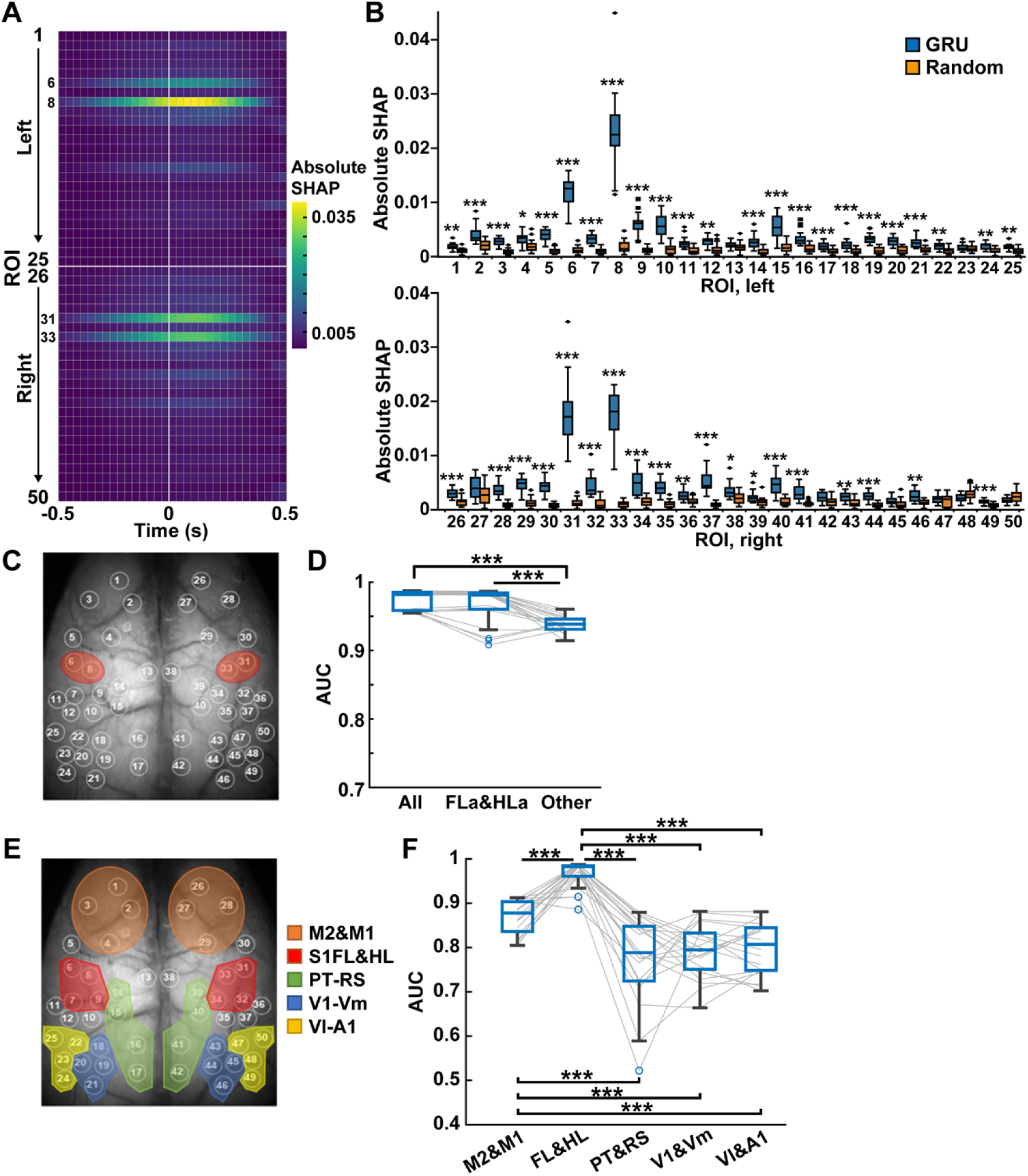
The forelimb and hindlimb areas of the somatosensory cortex contribute to behavioral state classification. (A) The absolute SHAP values at each ROI during the input window across all GRU decoders (50 ROIs × 31 frames (−0.5 ∼ 0.5 s) on 20 models average). (B) The absolute SHAP values for all frames at each ROI in GRU decoders with preprocessing data (GRU) and randomly shuffled data (Random). \**P* < 0.05, \*\**P* < 0.01, \*\*\**P* < 0.001, Wilcoxon rank-sum test with Holm correction, n = 20 models. See S2 Fig for ROIs 1–50. (C) Red ovals indicate the position of the somatosensory cortex anterior forelimb and hindlimb areas (ROIs 6, 8, 31, and 33). (D) Decoder performance using fluorescent signals from all cortical areas (All), somatosensory cortex anterior forelimb and hindlimb areas (FLa&HLa, ROIs 6, 8, 31, and 33), and the other 46 ROIs (Other). \*\*\**P* < 0.001, Wilcoxon rank-sum test with Holm correction, n = 20 models. (E) The ROIs were divided into five parts: motor areas (M2&M1, ROIs 1–4 and 26–29), somatosensory limb areas (FL&HL, ROIs 6–9 and 31–34), parietal and retrosplenial areas (PT&RS, ROIs 14–17 and 39–42), primary visual and visual medial areas (V1&Vm, ROIs 18–21 and 43–46), and visual lateral and auditory area (Vl&A1, ROIs 22–25 and 47–50). (F) Decoder performance using fluorescent signals from M2&M1, FL&HL, PT&RS, V1&Vm, and Vl&A1. \*\*\**P* < 0.001, Wilcoxon rank-sum test with Holm correction, n = 20 models.

Based on the results of SHAP, we trained the model using input data only from FLa and HLa (ROIs 6, 8, 31, and 33) and confirmed the performance of the behavioral classification (**Fig 6C**). We masked the signals out of these areas by replacing them with value 0 and used the masked data to train and test the GRU decoder (FLa&HLa). Oppositely, we masked the signals in FLa and HLa with 0 and trained and tested the GRU decoder (Other). The decoder performance using the somatosensory areas was compatible with the decoder trained with all area data (FLa&HLa, 0.966 ± 0.026; mean ± SD, n = 20 models; **Fig 6D**). However, the decoder using other cortical areas underperformed (Other, 0.938 ± 0.011; mean ± SD, n = 20 models; **Fig 6D**).

We further tested the group of cortical areas. We divided bilateral cortical areas into five parts (motor areas (M2&M1, ROIs 1–4, 26–29); somatosensory limb areas (FL&HL, ROIs 6–9, 31–34); parietal and retrosplenial areas (PT&RS, ROIs 14–17, 49– 52); primary visual and medial visual areas (V1&Vm, ROIs 18–21, 43–46); lateral visual and auditory areas (Vl&A1, ROIs 22–25, 47–50); **Fig 6E**) and used them separately for GRU training. The decoder performances were 0.869 ± 0.037 in M2&M1, 0.966 ± 0.030 in FL&HL, 0.776 ± 0.097 in PT&RS, 0.793 ± 0.060 in V1&Vm, and 0.798 ± 0.058 in Vl&A1 (mean ± SD, n = 20 models, respectively; **Fig 6F**). Consistent with the results in Fig. 5B, the decoder trained with FL&HL classified behavioral states with the highest accuracy. Moreover, the motor area’s decoder outperformed other cortical areas except for FL&HL. The correlation of the cortical activities with dynamics of behavioral states was weakly positive in all areas (mean ± SD; 0.21 ± 0.10, n = 50 ROIs; **S4 Fig**), which could not explain the predominance of the somatosensory limb areas in the GRU decoders.

In summary, our methods accurately classified mouse behavioral states from cortex-wide functional images consistent across mice and identified the essential features of cortical areas for behavioral classification in deep learning with both CNN and RNN. These results suggest the possibility of generalized neural decoding of voluntary behaviors with a whole-body motion from the cortical activity and the generation of interpretable decoders by end-to-end approach.

## Discussion

### Advantages of end-to-end behavior decoding from cortical calcium imaging

The present study demonstrated that deep learning using CNN-based end-to-end approaches accurately decoded the mouse behavioral states from cortical activity measured by mesoscopic calcium imaging. Recently, attempted speech and handwriting movements have been decoded on the temporal scales in real-time from the cortical activity obtained by microelectrode array and electrocorticography (ECoG) from human patients (Makin et al., 2020; Pan et al., 2018; Willett et al., 2021). Compared with the electrical recordings, calcium imaging is temporally slow but spatially high with a variable range of resolution from synaptic and cellular to regional scales. In CNN-RNN decoders, the robust performance of behavior classification was obtained using an input window from 0.067 s to 0.5 s. Our results indicate that the high spatial resolution of the calcium imaging contains sufficient information for decoding the mouse behavior even in the sub-second temporal order.

Furthermore, we visualized the importance of brain areas, the somatosensory cortex limb areas, for behavioral classification by the CNN-based end-to-end approach. These areas were commonly detected in the CNN-RNN decoders, suggesting that models were generalized between mice. Regional cortical activity in the somatosensory areas contributed to the decoding performance, supported by the RNN decoders. Since mice receive sensory inputs from the left and right limbs when moving on and touching the treadmill, the regional activity in the somatosensory areas may be reflected as a featured cortical response during locomotion. In addition, the primary somatosensory cortex also receives prior information about future movements from the primary motor cortex (Umeda et al., 2019). Utilizing the neural information from input-output relationships, such as the motor and somatosensory cortices, improves the performance of robotic arm control (Flesher et al., 2021). Our interpretable approach for deep learning decoders may help to identify multiregional cortical activities related to behavioral expressions.

### Combination of CNN and RNN for behavior decoding

Recently, a convolutional and recurrent neural network model has been applied to decoding finger trajectory from ECoG data, in which CNN was used to extract the features, and LSTM was used to capture the temporal dynamics of the signal (Xie et al., 2018). Similar to this architecture, our decoder with CNN-RNN effectively worked for mouse behavior classification and was superior to the decoder with CNN alone. Furthermore, the architecture LSTM followed by CNN was also applied to decoding the brain activity using EEG by reconstructing the visual stimuli, and it performed more accurately than the architecture CNN followed by LSTM (Zheng et al., 2020). The direction of architectures should be considered as a critical factor in the case of the combination of deep learning methods. By expanding the application of these methods in neuroscience research, behavior decoding from brain activity can deal with more complex patterns of behaviors with high temporal information, leading to the further development of BCI technologies.

## Materials and Methods

### Datasets

We used the previously reported dataset, including the 18,000-frame images of fluorescent signals in the cortex measured by mesoscopic calcium imaging at 30 frames/second and the time-matched behavioral states of locomotion and rest from head-fixed mice (Nakai et al., 2023). The dataset contains 64 sessions (for 10 min/session) from five Emx1G6 mice. The number of sessions in each mouse was 11, 12, 14, 15, and 12. We used all images (128 × 128 pixels × 18,000 frames × 64 sessions) for deep learning decoding with CNN and RNN. For deep learning analysis, we divided the five mice into subgroups at the rate of 3:1:1 for training, validation, and testing, respectively, to perform cross-validation, generating the twenty models in total (four models for each testing). For behavioral labeling, the frames with a locomotion speed more significant or less than 0.5 cm/s were defined as a state of “Run” or “Rest,” respectively.

### Data analysis

#### Deep learning with CNN-RNN

Deep learning with CNN-RNN was performed using Python 3.6, Anaconda Packages, PyTorch (https://pytorch.org), and fastai (https://docs.fast.ai). We used a PC equipped with Ubuntu 18.04 OS and NVIDIA GeForce RTX3090 GPU. All images were normalized by subtracting the average intensity in each pixel. The normalized images were divided by the variance of intensities of all pixels. For CNN classification, all images were then converted to an RGB image *I_t_* by combining three consecutive images from one frame before (red, *t* −1) to one frame after (blue, *t* +1) the target image *t* (green) with labeling a behavioral state of the target image *t* (Fig 2A). As the architecture of CNN, EfficientNet B0 was used from the Python package in GitHub (https://github.com/lukemelas/EfficientNet-PyTorch) (Tan and Le, 2020).

First, we trained the CNN to classify the behavioral state from the RGB images in the same manner of data allocation as deep learning with RNN. For the initial values of the CNN, we used the publicly available model that was pre-trained by ImageNet (Russakovsky et al., 2015). We used the random batches of size 512 using Adam optimizer (https://keras.io/api/optimizers/adam/ (Kingma and Ba, 2017)), binary cross-entropy loss function, and one-cycle training with a maximum learning rate of 0.001. After training, 1,280 features were extracted and fully connected to an output node. The activation function of the output node was set as sigmoid for binary classification of behavior labels. The number of epochs was set to 3. The model with the lowest loss in the validation data was adopted.

Next, a two-step training with CNN and RNN was performed for behavior state classification. Following the CNN training (Step 1), in which the initial values were set to the CNN models trained at the first stage, the RNN was trained using input data of sequential RGB images (Step 2). The inputs of RGB images for CNN were initially eleven consecutive images ranging from 0.17 s before (*I_t_* −5) to 0.17 s after (*I_t_* +5) the image *t,* which was labeled with the behavioral state at image *I_t_*(Fig 2A). After the convolution layer of CNN, 1,280 features per image were extracted by compression with average pooling and recursively input to RNN. The GRU and LSTM were used as the RNN architectures, which consisted of 128 units, 2 layers, and a dropout of 0.2. The hyperbolic tangent function was used as an activation function for RNN. The RNN units in the second layer were then fully connected to an output node. The activation function of the output node was set to sigmoid for the binary classification of behavior labels. We used the random batches of size 32 using Adam optimizer, binary cross-entropy loss function, and one-cycle training with the maximum learning rate of 0.001. The number of epochs was set to 3. The mixed precision (https://docs.fast.ai/callback.fp16.html) was used to improve the efficiency of the two-step training. We evaluated the loss for each Epoch and adopted the model with the lowest loss in the validation data. To compare the size of the input data for the CNN-RNN classification, we tested four different lengths of the time window, i.e., 0.067 s (*t* ±2), 0.17 s (*t* ±5), 0.33 s (*t* ±10), and 0.5 s (*t* ±15) before and after the image *t* (Fig 2F). The decoder performance was evaluated by the area under the receiver operating characteristic curve (AUC) for the classification of the test data.

#### Cut-out importance

We quantified the critical areas of images which contributed to the behavioral classification in the CNN-RNN decoder. The image (128 × 128 pixels) was divided into a 32-pixel square with a 16-pixel overlap, and each end was connected to the opposite end, thus obtaining 64 compartments. Before the CNN-RNN training, all pixels in a compartment were masked with a value of 0. We then trained the CNN-RNN by excluding information in the masked compartment area. Each compartment was scored by importance score, calculated by subtracting the AUC using the decoder trained with the masked data from the AUC using the decoder with the unmasked data.

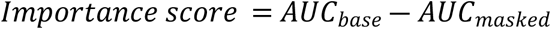

The importance score indicates how much the decoder performance using masked data (AUC_masked_) decreased compared to unmasked data (AUC_base_). The importance scores at one-fourth of the 32-pixel square were averaged among four times overlaps at the different masked areas and plotted on an 8×8 heat map. Then, the heat maps were averaged across all models. We named this analysis “cut-out importance.”

#### Preprocessing of regional cortical activity

This analysis was performed using MATLAB (MathWorks). The changes in cortical activity were calculated from fluorescent signals at the 50 regions of interest (ROIs) in the cortex (25 ROIs in each hemisphere), which was represented by dF/F, a percentage of changes from the baseline fluorescence (Nakai et al., 2023). In this study, a 1,000-frame moving average of dF/F was subtracted from dF/F to attenuate baseline variation of the fluorescent changes, which was an optimal filter size (S3 Fig).

#### Deep learning with RNN

Deep learning with recurrent neural network (RNN) was performed using Python 3.6 (https://www.python.org/), Anaconda Packages (https://docs.anaconda.com/anaconda/packages/old-pkg-lists/2021.05/py3.6_win-64/), TensorFlow (https://www.tensorflow.org/) and Keras (https://keras.io/). A PC with Ubuntu 16.04 OS and NVIDIA GeForce RTX2080 GPU was used. The code for deep learning is available in the following GitHub repository (https://github.com/atakehiro/Neural_Decoding_from_Calcium_Imaging_Data).

For binary classification of behavioral states, we assigned a value of 1 and 0 to the frames labeled “Run” and “Rest,” respectively. The input data of deep learning was 31 frames of the preprocessed dF/F, which localized from 15 frames before to 15 frames after a behavior-labeled frame, and a one-frame sliding window was used to cover all except for the first and last 15 frames. This period ranged up to 0.5 s after the behavioral expression had been used in the previous study (Pan et al., 2018). Each input data was normalized by Min-Max Scaling. We used six RNN architectures of deep learning (simple RNN, LSTM, GRU, and their bidirectional counterparts) for behavior classification in the same manner. The deep learning was trained with the random batches of size 256 using Adam optimizer (Kingma and Ba, 2017) and binary cross-entropy loss function. The unit number of RNN was set to 32. The hyperbolic tangent function was used as an activation function. The RNN is followed by a one-node fully connected layer. The activation function of the last classification node was set to sigmoid for the binary classification of behavior labels, and the label smoothing was set to 0.01. The number of epochs was set to 30, in which the models reached a stable loss and accuracy for the training and validation data. The model in the epoch with the lowest loss in the validation data was adopted. As a control, we generated the models trained with the behavioral labels permuted randomly (Random) and the models trained with non-preprocessed dF/F (Raw). The decoder performance was evaluated by the AUC for the classification of the test data.

#### Analysis of temporal differences in the input window using RNN decoders

To investigate the optimal conditions, we compared GRU decoders trained using the different lengths of the input time window and the temporally shifted target labels of behavioral classification (Fig 5). The target labels have temporally shifted the position from the center of the time window in the ranges from −2 to 2 s (from −60 to 60 frames) at 10-frames steps. The lengths of time window size 5, 10, 15, 20, 25, and 30, and the shifts of target label −60, −50, −40, −30, −20, −10, 0, 10, 20, 30, 40, 50, and 60 were analyzed.

#### Deep SHAP

We used Deep SHAP (the SHAP Python package in GitHub (https://github.com/slundberg/shap)) to visualize the basis for deep learning classifications. Deep SHAP is one of the feature attribution methods designed by combining SHAP (SHapley Additive exPlanation), which assigns each feature an importance value for machine learning predictions, with DeepLIFT, which is an additive feature attribution method that satisfies local accuracy and missingness (Lundberg and Lee, 2017). In this analysis, we randomly selected 10,000 frames from the test data (total 198,000-270,000 frames/test) to calculate SHAP values of each ROI, indicating the extent of contribution to the model output. The absolute SHAP values were averaged and represented as the overall importance of each ROI.

### Statistics

All statistical analyses were conducted in MATLAB (MathWorks). All bar plots with error bars represent mean ± SD. All box plots represent the median with interquartile range (IQR) (box) and 1.5 × IQR (whiskers), gray lines indicate the line plot of individual results, and ’o’ symbols indicate the outlier. For all statistical tests, the normality of the data and equal variance of groups were not assumed, and non-parametric tests were used for group comparisons. Wilcoxon rank-sum test with Holm correction was used. The significance level was set to *P* < 0.05.

## Acknowledgment

This work was supported in part by KAKENHI from JSPS, JP16H06316, JP16H06463, JP21H00202, JP21H04813, and JP21K19351; Japan Agency for Medical Research and Development (JP21wm0425011); Japan Science and Technology Agency (JPMJMS2299, JPMJMS229B); Intramural Research Grant (30-9) for Neurological and Psychiatric Disorders of NCNP; The Takeda Science Foundation, Research Foundation for Opto-Science and Technology, Taiju Life Social Welfare Foundation, The Naito Foundation, and The Tokumori Yasumoto Memorial Trust for Researches on Tuberous Sclerosis Complex and Related Rare Neurological Diseases. NN was also supported by KAKENHI from JSPS: JP19H04942.

## Author contributions

Conceptualization, TA and NN; methodology, TA, NN, and OY; investigation, TA; visualization, TA and NN; supervision, NN and TT; writing—original draft, TA; writing—review & editing: NN, OY, and TT; funding acquisition, NN and TT.

## Competing interests

The authors have no conflict of interest.

## Data and materials availability

All data are available by the authors upon reasonable request. Codes are available here: https://github.com/atakehiro/Neural_Decoding_from_Calcium_Imaging_Data.

## Supporting information

S1 Fig. Importance scores in each session.

S2 Fig. Fluorescent calcium signals and corresponding cortical areas.

S3 Fig. Preprocessing of the fluorescent signals for deep learning classification. S4 Fig. Correlation between fluorescent signals and locomotor activity.

## Notes

### Competing Interest Statement

The authors have declared no competing interest.

## References

Chang, C.-J., Guo, W., Zhang, J., Newman, J., Sun, S.-H., Wilson, M., 2021. Behavioral clusters revealed by end-to-end decoding from microendoscopic imaging. https://doi.org/10.1101/2021.04.15.440055

Cho, K., van Merrienboer, B., Gulcehre, C., Bahdanau, D., Bougares, F., Schwenk, H., Bengio, Y., 2014. Learning Phrase Representations using RNN Encoder-Decoder for Statistical Machine Translation. https://doi.org/10.48550/arXiv.1406.1078

Craik, A., He, Y., Contreras-Vidal, J.L., 2019. Deep learning for electroencephalogram (EEG) classification tasks: a review. J. Neural Eng. 16, 031001. https://doi.org/10.1088/1741-2552/ab0ab5

Etter, G., Manseau, F., Williams, S., 2020. A Probabilistic Framework for Decoding Behavior From in vivo Calcium Imaging Data. Front Neural Circuits 14, 19. https://doi.org/10.3389/fncir.2020.00019

Flesher, S.N., Downey, J.E., Weiss, J.M., Hughes, C.L., Herrera, A.J., Tyler-Kabara, E.C., Boninger, M.L., Collinger, J.L., Gaunt, R.A., 2021. A brain-computer interface that evokes tactile sensations improves robotic arm control. Science 372, 831–836. https://doi.org/10.1126/science.abd0380

Gunning, D., Stefik, M., Choi, J., Miller, T., Stumpf, S., Yang, G.-Z., 2019. XAI-Explainable artificial intelligence. Sci Robot 4, eaay7120. https://doi.org/10.1126/scirobotics.aay7120

Hochberg, L.R., Bacher, D., Jarosiewicz, B., Masse, N.Y., Simeral, J.D., Vogel, J., Haddadin, S., Liu, J., Cash, S.S., van der Smagt, P., Donoghue, J.P., 2012. Reach and grasp by people with tetraplegia using a neurally controlled robotic arm. Nature 485, 372–375. https://doi.org/10.1038/nature11076

Hochreiter, S., Schmidhuber, J., 1997. Long short-term memory. Neural Comput 9, 1735– 1780. https://doi.org/10.1162/neco.1997.9.8.1735

Kingma, D.P., Ba, J., 2017. Adam: A Method for Stochastic Optimization. https://doi.org/10.48550/arXiv.1412.6980

LeCun, Y., Bengio, Y., Hinton, G., 2015. Deep learning. Nature 521, 436–444. https://doi.org/10.1038/nature14539

Li, C., Chan, D.C.W., Yang, X., Ke, Y., Yung, W.-H., 2019. Prediction of Forelimb Reach Results From Motor Cortex Activities Based on Calcium Imaging and Deep Learning. Front Cell Neurosci 13, 88. https://doi.org/10.3389/fncel.2019.00088

Livezey, J.A., Glaser, J.I., 2021. Deep learning approaches for neural decoding across architectures and recording modalities. Brief Bioinform 22, 1577–1591. https://doi.org/10.1093/bib/bbaa355

Lundberg, S., Lee, S.-I., 2017. A Unified Approach to Interpreting Model Predictions. https://doi.org/10.48550/arXiv.1705.07874

Makin, J.G., Moses, D.A., Chang, E.F., 2020. Machine translation of cortical activity to text with an encoder-decoder framework. Nat Neurosci 23, 575–582. https://doi.org/10.1038/s41593-020-0608-8

Murano, T., Nakajima, R., Nakao, A., Hirata, N., Amemori, S., Murakami, A., Kamitani, Y., Yamamoto, J., Miyakawa, T., 2022. Multiple types of navigational information are independently encoded in the population activities of the dentate gyrus neurons. Proc Natl Acad Sci U S A 119, e2106830119. https://doi.org/10.1073/pnas.2106830119

Nakai, N., Sato, M., Yamashita, O., Sekine, Y., Fu, X., Nakai, J., Zalesky, A., Takumi, T., 2023. Virtual reality-based real-time imaging reveals abnormal cortical dynamics during behavioral transitions in a mouse model of autism. Cell Rep 112258. https://doi.org/10.1016/j.celrep.2023.112258

Pan, G., Li, J.-J., Qi, Y., Yu, H., Zhu, J.-M., Zheng, X.-X., Wang, Y.-M., Zhang, S.-M., 2018. Rapid Decoding of Hand Gestures in Electrocorticography Using Recurrent Neural Networks. Front Neurosci 12, 555. https://doi.org/10.3389/fnins.2018.00555

Ren, C., Komiyama, T., 2021. Characterizing Cortex-Wide Dynamics with Wide-Field Calcium Imaging. J. Neurosci. 41, 4160–4168. https://doi.org/10.1523/JNEUROSCI.3003-20.2021

Russakovsky, O., Deng, J., Su, H., Krause, J., Satheesh, S., Ma, S., Huang, Z., Karpathy, A., Khosla, A., Bernstein, M., Berg, A.C., Fei-Fei, L., 2015. ImageNet Large Scale Visual Recognition Challenge. Int J Comput Vis 115, 211–252. https://doi.org/10.1007/s11263-015-0816-y

Schwemmer, M.A., Skomrock, N.D., Sederberg, P.B., Ting, J.E., Sharma, G., Bockbrader, M.A., Friedenberg, D.A., 2018. Meeting brain–computer interface user performance expectations using a deep neural network decoding framework. Nat Med 24, 1669–1676. https://doi.org/10.1038/s41591-018-0171-y

Skomrock, N.D., Schwemmer, M.A., Ting, J.E., Trivedi, H.R., Sharma, G., Bockbrader, M.A., Friedenberg, D.A., 2018. A Characterization of Brain-Computer Interface Performance Trade-Offs Using Support Vector Machines and Deep Neural Networks to Decode Movement Intent. Front Neurosci 12, 763. https://doi.org/10.3389/fnins.2018.00763

Tan, M., Le, Q.V., 2020. EfficientNet: Rethinking Model Scaling for Convolutional Neural Networks. https://doi.org/10.48550/arXiv.1905.11946

Umeda, T., Isa, T., Nishimura, Y., 2019. The somatosensory cortex receives information about motor output. Sci Adv 5, eaaw5388. https://doi.org/10.1126/sciadv.aaw5388

Vega García, M., Aznarte, J.L., 2020. Shapley additive explanations for NO2 forecasting. Ecological Informatics 56, 101039. https://doi.org/10.1016/j.ecoinf.2019.101039

Willett, F.R., Avansino, D.T., Hochberg, L.R., Henderson, J.M., Shenoy, K.V., 2021. High-performance brain-to-text communication via handwriting. Nature 593, 249–254. https://doi.org/10.1038/s41586-021-03506-2

Xie, Z., Schwartz, O., Prasad, A., 2018. Decoding of finger trajectory from ECoG using deep learning. J Neural Eng 15, 036009. https://doi.org/10.1088/1741-2552/aa9dbe

Zheng, X., Chen, W., Li, M., Zhang, T., You, Y., Jiang, Y., 2020. Decoding human brain activity with deep learning. Biomedical Signal Processing and Control 56, 101730. https://doi.org/10.1016/j.bspc.2019.101730

